# Cdc42 GTPase regulates ESCRTs in nuclear envelope sealing and ER remodeling

**DOI:** 10.1101/808436

**Authors:** Michelle S. Lu, David G. Drubin

## Abstract

Small GTPases of the Rho family are binary molecular switches that regulate a variety of processes including cell migration and oriented cell divisions. Known Cdc42 effectors include proteins involved in cytoskeletal remodeling and kinase-dependent transcription induction, but none involved in the maintenance of nuclear envelope integrity or endoplasmic reticulum (ER) morphology. Maintenance of nuclear envelope integrity requires the EndoSomal Complexes Required for Transport (ESCRT) proteins, but how they are regulated in this process remains unknown. Here we show by live-cell imaging a novel Cdc42 localization with ESCRT proteins at sites of nuclear envelope and ER fission, and by genetic analysis, uncover a unique Cdc42 function in regulation of ESCRT proteins at the nuclear envelope and sites of ER tubule fission. Our findings implicate Cdc42 in nuclear envelope sealing and ER remodeling, where it regulates ESCRT disassembly to maintain nuclear envelope integrity and proper ER architecture.

**Summary:** The small Rho GTPase Cdc42 is a well-known regulator of cytoskeletal rearrangement and polarity development in all eukaryotic cell types. Here, Lu and Drubin report the serendipitous discovery of a novel Cdc42-ESCRT-nuclear envelope/endoplasmic reticulum connection.

## Introduction

Maintenance of the nuclear envelope is critical for compartmentalization of the activities of the nucleus away from those of the cytoplasm, and requires the proper function of several proteins including the ESCRT and LEM-family proteins (Gu et al., 2017; Olmos et al., 2016; Olmos et al., 2015; Thaller et al., 2019; Vietri et al., 2015; Webster et al., 2014; Webster et al., 2016). The ESCRTs are a family of membrane-deforming proteins that mediate reverse-topology scission of membranes (Schöneberg et al., 2016). Together with the INM-embedded LEM-family proteins, the ESCRTs carry out the closure of holes (annular fusion) in the nuclear envelope (Olmos et al., 2016). Loss of proper ESCRT and LEM protein function in open-mitosis systems results in the failure of post-mitotic nuclear envelope reassembly, ultimately leading to delayed microtubule assembly, compromised nuclear integrity, and DNA damage (Olmos et al., 2015; Olmos et al., 2016; Vietri et al., 2015). In closed-mitosis systems, wherein the nuclear envelope does not breakdown and instead undergoes a fission process, ESCRT and LEM dysfunction result in the loss of nucleo-cytoplasmic partitioning (Gu et al., 2017; Thaller et al., 2019; Webster et al., 2014; Webster et al., 2016).

Proper ESCRT function relies on their ordered recruitment and assembly into organized filaments at membrane holes and areas of high membrane curvature (Schöneberg et al., 2016). The dysregulation of their assembly can result in toxic nuclear envelope malformations, such as those seen in fission yeast overexpressing Lem2p, an inner nuclear membrane LEM-family protein that recruits the ESCRT-III component Cmp7p in fission yeast (Gonzalez et al., 2012; Gu et al., 2017; King et al., 2006). Their disassembly must also be tightly regulated, as unrestricted ESCRT function can lead to large gaps in the nuclear envelope, as seen in fission yeast lacking Vps4, a type 1 AAA+ ATPase whose main cellular function appears to be to disassemble ESCRT filaments (Gu et al., 2017). Thus, both assembly and disassembly of ESCRTs must be carefully controlled *in vivo* to ensure nuclear envelope integrity.

While it is known that the ESCRT components, LEM-family proteins, and VPS4 are required for nuclear envelope sealing in mammalian cells, fission yeast, and budding yeast, how these components are regulated during this process is not yet clear. Here we report our serendipitous discovery of a novel Cdc42-ESCRT-nuclear envelope connection through live-cell imaging approaches, which, combined with biochemical and genetic analyses, reveal that the Rho family GTPase Cdc42 functions in regulation of ESCRT-III during nuclear envelope sealing and ER remodeling in budding yeast. Our data suggest that Cdc42 is a regulator of ESCRT filament disassembly, possibly mediating the union of ESCRT filaments and Vps4 at sites of annular fusion on the nuclear envelope and also to sites of ER tubule fission. This function represents a novel function of Cdc42, a molecular switch with well-known functions in actin polymerization (Kim et al., 2000; Prehoda et al., 2000), spindle positioning (Garrard et al., 2003; Gotta et al., 2001; Kay et al., 2001), and exocytosis (Adamo et al., 2001; Zhang et al., 2001).

## Results and Discussion

Functional Cdc42 has never been imaged in live budding yeast cells because it is not amenable to N- or C-terminal protein fusions. In an effort to visualize Cdc42 in living cells, we created a functional Cdc42-mCherry internal fusion protein (Cdc42-mCherry^sw^) in budding yeast analogous to that engineered previously in fission yeast (Bendezu et al., 2015). Unlike N-terminally tagged alleles of *CDC42* (Wu et al., 2015), expression of this internally tagged *CDC42* allele from its native promoter as the sole source of Cdc42 does not cause temperature sensitivity and cells are fully viable at 37 °C, though not at 39 °C (Fig. S1 A). Live-cell imaging of this strain reveals the expected localization of Cdc42 at sites of polarized growth (Adams et al., 1990; Etienne-Manneville, 2004) (Fig. 1 A). However, in addition to its expected localization, Cdc42-mCherry^sw^ showed a novel subcellular localization in 23 ± 6.9% of vegetatively growing cells (Fig. 1 B). These cells contained a single Cdc42 spot in the mother and/or the bud, and sometimes the spot segregated from the mother to the bud (Fig. S1 B, Video 1). In an effort to identify the function of this spot, we first surveyed subcellular structures with which the spot might be associated by examining strains coexpressing endogenously-tagged fluorescent organelle markers. We observed that the Cdc42 spot is associated with the vacuole, where it is either completely overlapping with, or juxtaposed to the vacuolar membrane, which we visualized using Vph1-GFP (Fig. S2 A). Additionally, we saw that the Cdc42 spot is associated with the ER (marked by GFP-HDEL), usually in an apposed rather than overlapping manner (Fig. S2 B). Having identified a new subcellular Cdc42 localization, we next wanted to determine if there is a function associated with the Cdc42 spot.

**Figure 1.**
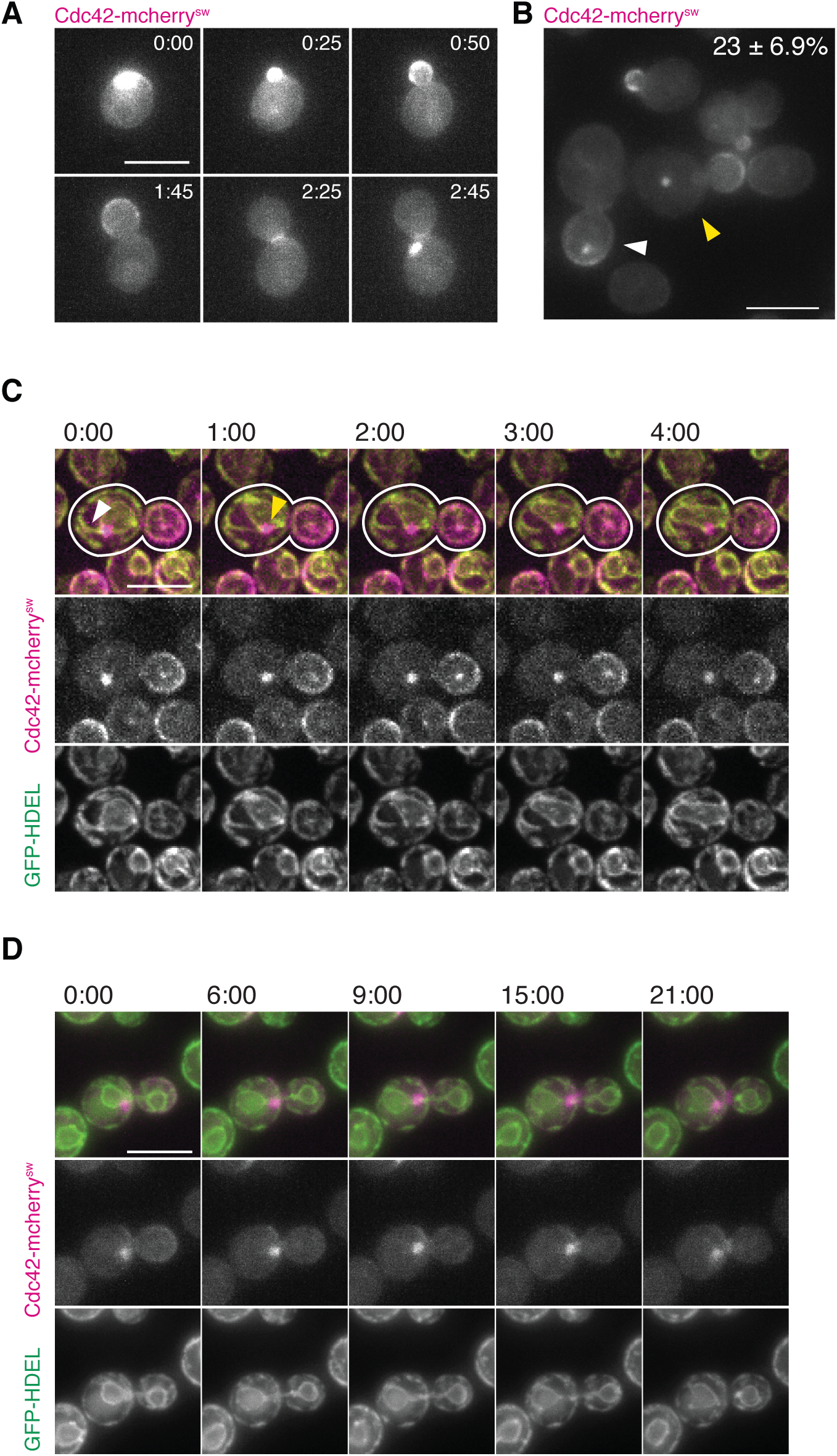
Cdc42-mCherry^SW^ localizes to a single spot at sites of ER remodeling in a subpopulation of cells. **(A)** Montage of wide-field epifluorescence time lapse imaging of cells endogenously expressing Cdc42-mCherry^SW^ during polarized cell growth. Cells were imaged in early log phase in minimal imaging media at 23-25 °C. 3µm maximum intensity projection at 0.5µm steps. All scale bars, 5µm. **(B)** Field of view of same strain in **A** showing two examples of the 23% ± 6.9% of cells with Cdc42 spots (± SD. n = 1527, four trials). White arrow indicates bud containing Cdc42 spot and yellow arrow indicates a budding mother containing Cdc42 spot. **(C)** Montage of confocal spinning disk movie stills of cells endogenously expressing Cdc42-mCherry^SW^ (magenta) and GFP-HDEL (green) undergoing fission of ER tubule emanating from the nucleus. White arrowhead in first frame indicates ER tubule and yellow arrowhead in second frame indicates site of severed ER tubule on the nuclear ER. 2µm maximum intensity projections at 0.5µm steps. **(D)** Montage of confocal spinning disk movie stills of the same cells as **A** undergoing nuclear fission. 3µm maximum intensity projections at 0.5µm steps.

Because Cdc42 had previously been implicated in vacuole function (Eitzen et al., 2001; Muller et al., 2001) but never in ER function, we decided to focus on the function of ER-associated Cdc42. We therefore analyzed the dynamic behavior of the ER-associated spot relative to the ER membrane. Live cell imaging revealed that the Cdc42 spot often appears at the base of ER tubules undergoing fission. Time lapse imaging of cells that exhibit fission of ER tubules emanating from the nucleus showed the Cdc42 spot localizing to the base of the ER tubule neck before, during, and after the fission event (Fig. 1 C, Video 2). We observed a similar behavior in cells undergoing nuclear fission late in mitosis, where, throughout the duration of nuclear fission, the Cdc42 spot remained near the base of the neck of the dividing nuclear envelope, whose outer membrane is continuous with the ER and thereby is marked by GFP-HDEL (Fig. 1 D, Video 3, Fig. S1 D).The Cdc42 spot not only appeared to be involved in nuclear and ER membrane remodeling, but also vacuolar membrane remodeling. The Cdc42 spot can be observed at the vertices of fusing vacuoles, remaining at the convergence points until the vacuole fragments have fully fused (Fig. S1 C, Video 4), lending further evidence of a potential membrane remodeling function for Cdc42. Having identified a novel subcellular localization of Cdc42 to organelles undergoing membrane remodeling, we next sought to determine if Cdc42 appears at these sites with proteins known to reshape intracellular membranes.

The ESCRT proteins have previously been found to remodel the same organelles that we found colocalized with Cdc42, so we tested whether Cdc42 and the ESCRTs were involved in the same membrane remodeling process. The ESCRT proteins are membrane deforming proteins known to function at endosomes and vacuoles (Zhu et al., 2017), and are also involved in the sealing of the nuclear envelope (Webster et al., 2016; Gu et al., 2017; Vietri et al., 2015; Olmos et al., 2016). We first examined the subcellular localization of Chm7, an ESCRT-III-like protein that functions in nuclear envelope sealing in fission and budding yeast (Webster EMBO, Gu), and whose mammalian homolog CHMP7 functions in nuclear envelope reformation and sealing in animal cells (Vietri et al., 2015; Olmos et al., 2016). As previously reported, we also found that Chm7 localizes to a single spot in a subpopulation of cells (Webster et al., 2016), and in cells harboring both Cdc42 and Chm7 foci, ∼70% of the Cdc42 spots colocalized with the Chm7 spot (Fig. 2 A). We also examined the subcellular localization of Snf7, an ESCRT-III protein that functions with Chm7 in the sequestration of defective nuclear pore complexes in budding yeast (Webster et al., 2014), and whose mammalian homolog CHMP4B is involved in nuclear envelope sealing in animal cells (Olmos et al., 2015). Unlike Chm7 and Cdc42, Snf7 formed a varying number of intracellular foci. Importantly, in cells containing a Cdc42 spot and at least one GFP-Snf7 spot, ∼70% of Cdc42 spots colocalized with a GFP-Snf7 spot (Fig. 2 B). To determine whether the Cdc42 spot with Snf7 is the same structure as the Cdc42 spot juxtaposed to the ER (Figs. 1 C, D, Fig. S2 B), we generated a strain expressing Cdc42-mcherry^SW^, GFP-HDEL, and mtagBFP2-Snf7 for three-color live-cell imaging. We observed that ER-associated Cdc42 spots also contained Snf7 (Fig. 2 C), suggesting that they are the same structure.

**Figure 2.**
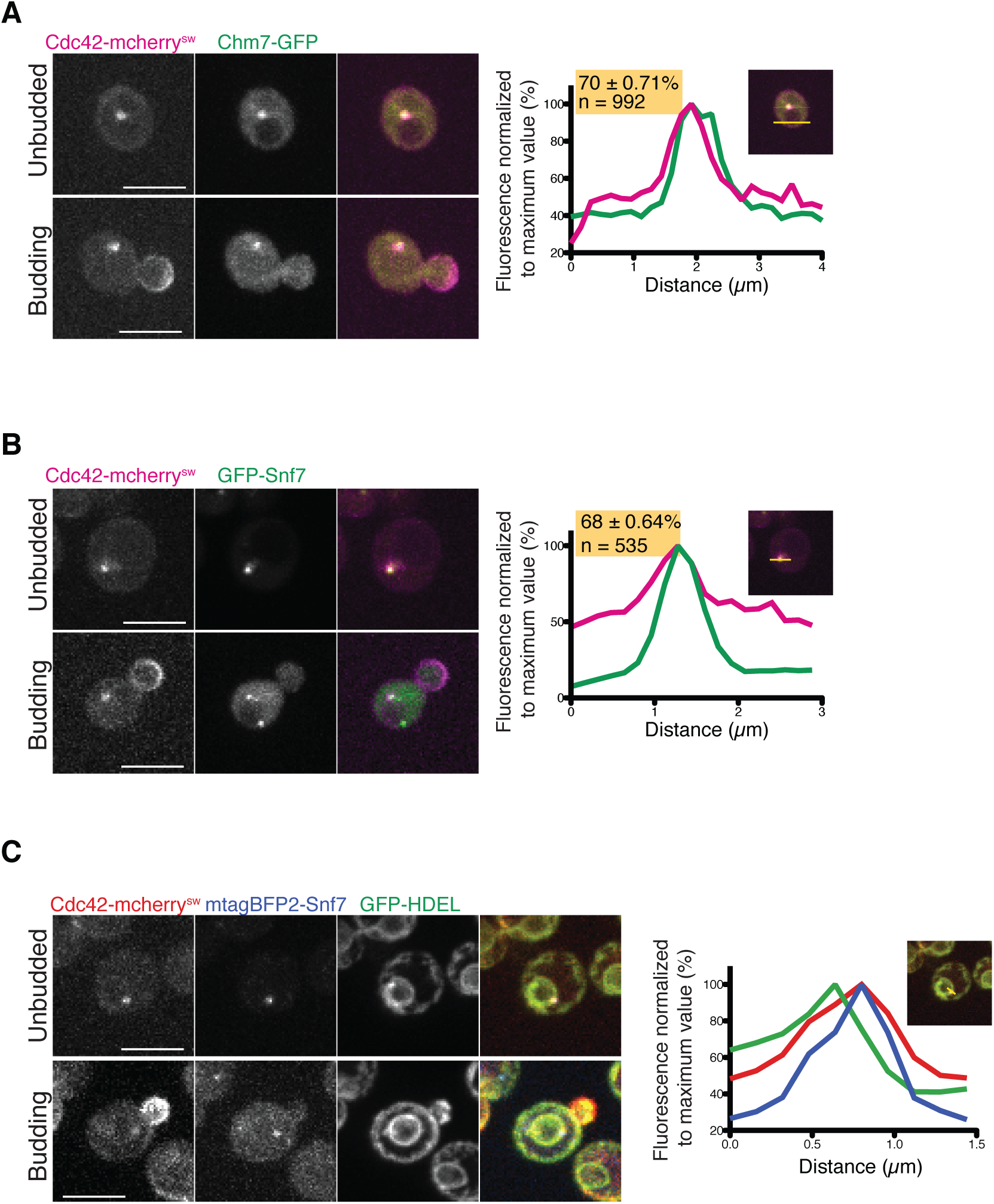
Cdc42 colocalizes with ESCRTs in vivo. **(A)** Confocal spinning disk still images of cells endogenously expressing Cdc42-mCherry^SW^ (magenta) and Chm7-GFP (green) in unbudded (top panel) or budded cell (lower panel). Plot of normalized fluorescence intensity profiles of Chm7-GFP (green) and Cdc42-mCherry^SW^ (magenta) along yellow line shown in inset image. Cells that contain both Cdc42 spot and Chm7 spot where both are colocalized = 70 ± 0.71% (SD. n = 992, two trials). **(B)** Confocal spinning disk stills of cells endogenously expressing Cdc42-mCherry^SW^ (magenta) and GFP-Snf7 (green) in unbudded (top panel) or budded cell (lower panel). Plot of normalized fluorescence intensity profiles of GFP-Snf7 (green) and Cdc42-mCherry^SW^ (magenta) along yellow line shown in inset image. Cells that contain both Cdc42 spot and Snf7 punctae, where they are colocalized = 68 ± 0.64% (SD. n = 535, two trials). **(C)** Confocal spinning disk stills of cells endogenously expressing Cdc42-mCherry^SW^ (red), GFP-HDEL (green), and mtagBFP2-Snf7 (blue) in G1 phase (top panel) or budding (lower panel). Plot of normalized fluorescence intensity profiles of Cdc42 (magenta), ER (green), and Snf7 (blue) along yellow line shown in inset image. All images are 2µm maximum intensity projections at 0.5µm steps. All scale bars 5µm.

Live imaging of Chm7 and Snf7 suggested that they were involved in a similar putative ER membrane remodeling process as Cdc42. Like the Cdc42 spot, we observed instances of the Chm7 spot and Snf7 punctae localizing to the ER bottleneck areas of nuclear envelope fission (Figs. 3 A, B). This localization behavior is consistent with numerous studies detailing ESCRT proteins’ preferences for high-curvature membranes, particularly necks and holes (Cashikar et al., 2014; Lee et al., 2015; De Franceschi et al., 2018; McCollough et al., 2015). Three-color live-cell imaging revealed that Cdc42 and Snf7 colocalize to the same peri-nuclear structure, that, in cells undergoing cytokinesis, appears to migrate to the base of an ER tubule undergoing nuclear fission (Fig. 3C, Video 5). These observations suggest that both Cdc42 and Snf7 function together at ER fission sites. These live-cell imaging observations point to a membrane-remodeling function for Cdc42, and together with the colocalization with ESCRT proteins, suggest that Cdc42 functions with Snf7 and Chm7 at the ER membrane.

**Figure 3.**
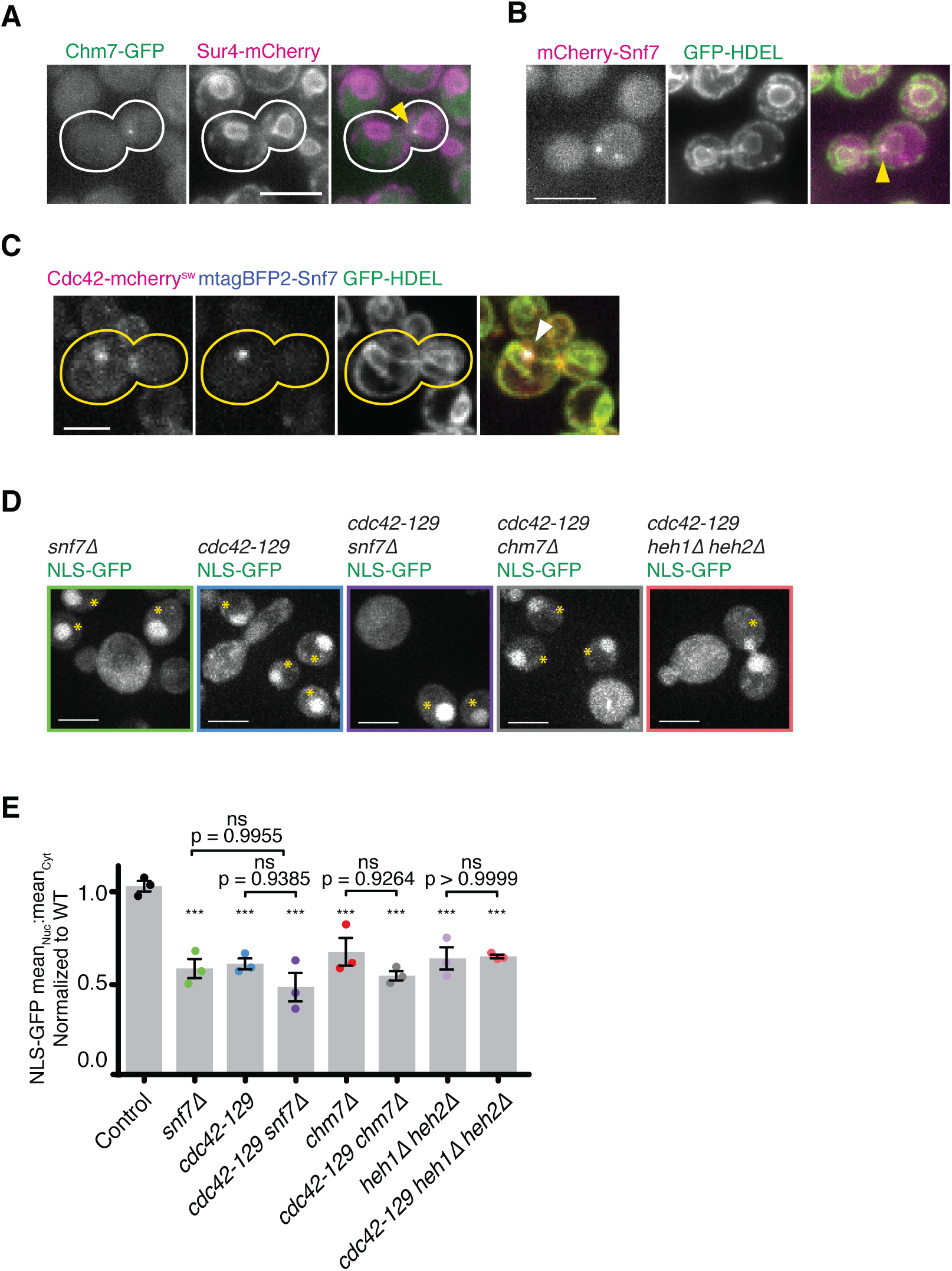
Cdc42 functions in the same NE sealing pathway as ESCRT and LEM proteins. **(A)** Stills from epifluorescence movies of cells endogenously expressing Chm7-GFP (green) and Sur4-mCherry (magenta), to mark the nuclear envelope. White arrowhead indicates Chm7 spot at post-fission nuclear membrane. **(B)** Stills from epifluorescence movies of cells endogenously expressing mCherry-Snf7 (magenta) and GFP-HDEL (green), to mark the ER. White arrowhead indicates Snf7 punctum at base of ER tubule neck undergoing nuclear fission. **(C)** Confocal spinning disk still of a cell endogenously expressing Cdc42-mCherry^SW^ (red), GFP-HDEL (green), mtagBFP2-Snf7 (blue) undergoing nuclear fission. White arrowhead indicates Snf7 and Cdc42 structure on ER membrane. **(D)** Confocal spinning disk micrographs of indicated mutant strains, with wild-type cells (marked with yellow asterisks) in the same field of view. **(E)** Quantification of each mutant’s mean nuclear:cytosolic GFP fluorescence intensity normalized to that of wild-type cells in the same field of view. Colored dots report means from individual trials. Asterisks denote statistical significance compared with control after one-way ANOVA (F = 10.74), followed by Sidak’s multiple comparison test to determine p-values between bracketed strains. n, means, and SDs of individual trials reported in supporting information. **A-C** are 2µm maximum intensity projections at 0.5µm steps. **D** is 11µm maximum intensity projection at 0.5µm steps. All cells in this figure were grown to log phase in minimal imaging media at 23-25 °C. All scale bars, 5µm.

Because ESCRT proteins have been extensively documented to function in nuclear envelope sealing in budding yeast, fission yeast, and mammalian cells (Webster et al., 2014; Webster et al., 2016; Gu et al., 2017’ Vietri et al., 2015; Olmos et al., 2015), we reasoned that Cdc42 may coordinate with Chm7 and Snf7 to seal the hole left behind in the nuclear envelope after ER tubule fission from the nuclear envelope. This possibility is consistent with findings from previous studies in which ESCRT-III depletion in mammalian cells disrupts the annular fusion of the reassembling nuclear envelope after mitosis, ultimately causing a reduction in the post-mitotic nucleo-cytoplasmic partitioning of a GFP-NLS probe (Vietri et al., 2015; Olmos et al., 2015). A similar leaky nucleus phenotype occurs in *chm7*Δ budding yeast, where the deletion of *CHM7* leads to a decrease in nuclear:cytosolic mean GFP-NLS fluorescence ratio at 37°C (Webster et al., 2016). To determine whether *CDC42* is also required for nuclear envelope sealing, we searched for *cdc42* mutants with a leaky nucleus phenotype and identified the *cdc42-129* allele (Kozminski et al., 2000). Using the GFP-NLS reporter assay, we found that the temperature-sensitive *cdc42-129* allele causes a leaky nucleus phenotype at the restrictive temperature of 37°C, phenocopying *chm7*Δ (Webster et al., 2016) (Figs. 3 D, E, Fig. S3). The *cdc42-129 chm7*Δ double mutant does not have an exacerbated phenotype, suggesting that these two proteins function in the same pathway. Similar to *chm7*Δ, *snf7*Δ cells also have leaky nuclei, and combining this deletion with *cdc42-129* gave the same phenotype as the single mutants (Figs. 3 D, E, Fig. S3). Thus, Cdc42 functions in the same pathway as the ESCRT-III proteins Snf7 and Chm7 in nuclear envelope sealing.

Further molecular genetic analysis revealed that Cdc42 functions with LEM family proteins in nuclear envelope resealing. Chm7 and Snf7 function with the LEM protein Heh2 and its paralog Heh1 in nuclear pore complex quality control and nuclear envelope sealing in budding yeast (Webster et al., 2014). In addition, in fission yeast and human cells, the LEM-family protein LEM2 recruits CHMP7 during nuclear envelope closure (Gu et al., 2017). To determine if Cdc42 is a component of this ESCRT-LEM protein complex that functions in nuclear envelope sealing, we tested for genetic interactions between mutant alleles of *CDC42* and *HEH1*/*HEH2*. As previously reported, we found that *heh1Δ heh2*Δ mutants have leaky nuclei (Webster et al., 2014) (Figs. 3 D, E, Fig. S3). Combining these mutations with the *cdc42-129* mutation did not result in a synthetic effect (Figs. 3 D, E, Fig. S3), suggesting that all three proteins function in the same pathway for this cellular function. Our genetic analyses suggest that *CDC42* functions in the same pathway with *HEH1*, *HEH2*, *SNF7*, and *CHM7* in nuclear envelope sealing. We next sought to elucidate the molecular mechanisms underlying Cdc42 function in this pathway.

How the ESCRT and LEM-family proteins are regulated during nuclear envelope sealing function is not yet known, so we investigated whether Cdc42 plays a regulatory role in this pathway. Many of the ESCRT-III proteins contain an auto-inhibitory regulatory C-terminal region whose de-repression leads to polymerization into active oligomers (Cashikar et al., 2014; Hanson et al., 2008; Henne et al., 2012; Shen et al., 2014; Tang et al., 2015). The disassembly of “open” ESCRT-III oligomers can be induced by the association with the type 1 AAA+ ATPase Vps4; the strength of the binding affinities of different ESCRT-III proteins for Vps4 are directly correlated with how well they are disassembled. While the mechanisms underlying Vps4-mediated disassembly of ESCRT-III proteins CHMP1 (Stuchell-Brereton et al., 2007), CHMP2 (Stuchell-Brereton et al., 2007; Obita et al., 2007), CHMP6 (Kieffer et al., 2008), and IST1 (Guo et al., 2015), which associate with each other tightly, are known, how Vps4 disassembles weakly- or non-interacting ESCRT-III proteins including CHMP4/Snf7, is not known, though it has been proposed that accessory proteins that bind both ESCRT-III and Vps4 might facilitate Vps4-mediated disassembly (Schöneberg et al., 2016).

To explore whether Cdc42 plays a direct or accessory role in Vps4-mediated ESCRT filament disassembly, we first determined whether Vps4 is present at sites of Cdc42’s nuclear function. Using a fully functional endogenous Vps4-3xHA-GFP (Adell et al., 2017), we observed that Vps4 punctae colocalize with the Cdc42 spot in 70% of cells harboring a Cdc42 spot (Fig. 4 A). Having established that Cdc42 colocalizes with the machinery required for ESCRT-III filament disassembly, we next determined if Cdc42 plays a role in Snf7 disassembly.

**Figure 4.**
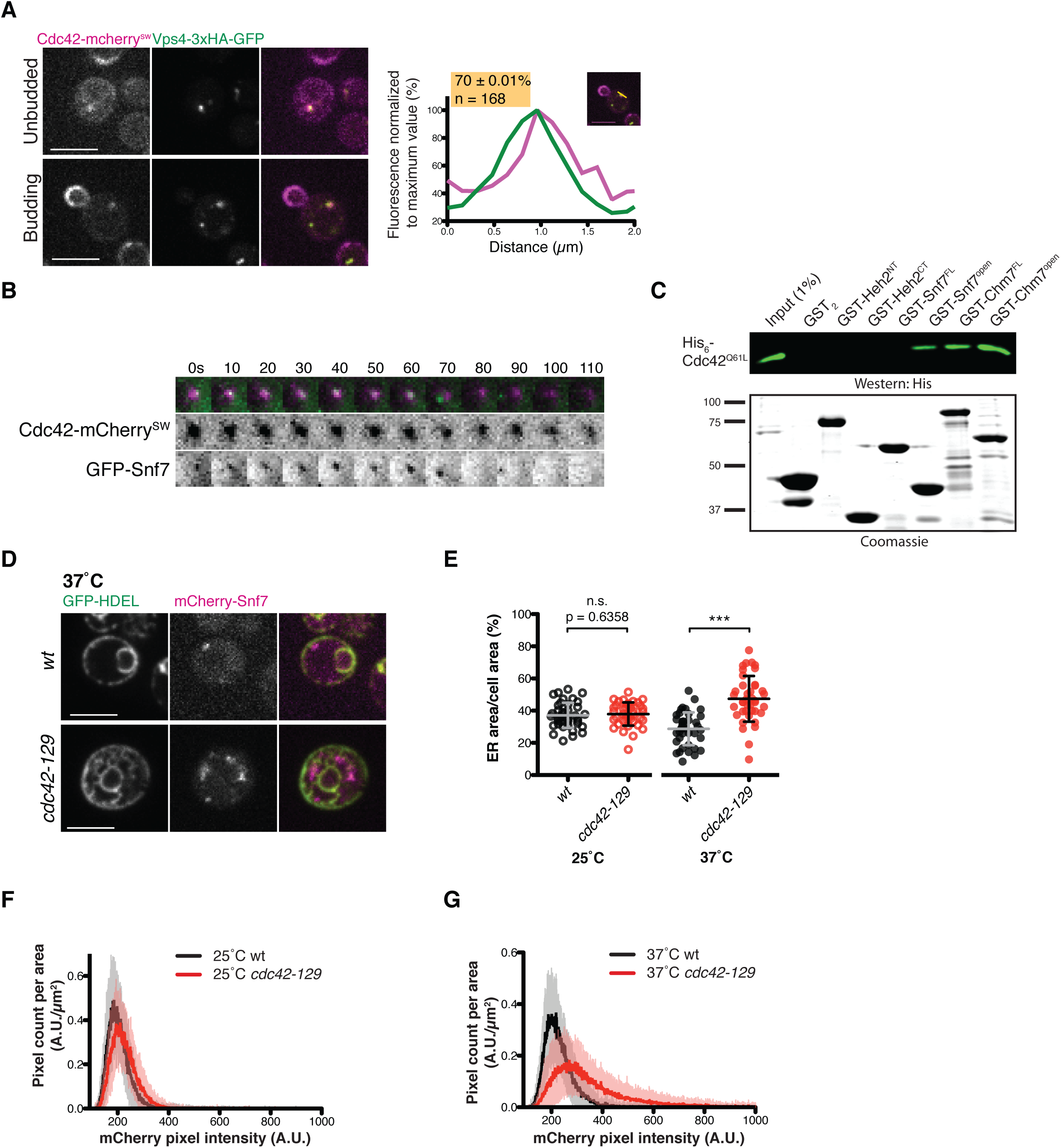
Cdc42 is at sites of Snf7 disassembly and its disruption leads to Snf7 aggregates and ER tubule extensions from the nuclear envelope. **(A)** Confocal spinning disk still images of cells endogenously expressing Cdc42-mCherry^SW^ (magenta) and Vps4-3xHA-GFP (green) in unbudded (top panel) or budded cell (lower panel). Plot of normalized fluorescence intensity profiles of Vps4-3xHA-GFP (green) and Cdc42-mCherry^SW^ (magenta) along yellow line shown in inset image. Cells that contain both Cdc42 spot and Vps4 spot where both are colocalized = 70.6 ± 0.01% (SD. n = 168). **(B)** Montage of stills from confocal spinning disk movie of a Cdc42 spot (magenta) and Snf7 (green) punctum. Time is in seconds. **(C)** GST-fusions of Heh2, Chm7, or Snf7 fragments or full-length protein (Coomassie, lower panel) were immobilized on GST-beads as bait and incubated with purified His_6_-Cdc42 (constitutively GTP-bound) as prey (detected by western blot, top panel). **(D)** Confocal spinning disk micrographs of wildtype (top panel) and *cdc42-129* (bottom panel) cells expressing GFP-HDEL (green) to mark ER and mCherry-Snf7 (magenta) grown to early-log phase in minimal media and shifted to 37°C for six hours. Single (medial) focal plane. All scale bars 5 µm. **(E)** Quantification of percentage cell area occupied by ER for same strains shown in **D**. Open circles indicate wildtype (red) and *cdc42-129* mutant strain (black) imaged at the permissive temperature (25°C), and closed circles at the restrictive temperature (37°C). Two trials, n = 40, mean ± SD. Asterisks denote statistical significance compared with control using student’s t-test. **(F)** Quantification of mCherry-Snf7 punctae of same strain shown in **D** grown to early-log phase in minimal media at 25°C, as a frequency distribution of the fluorescence intensity of each pixel in a cell normalized to cell area (µm^2^). Two trials, n = 40, mean ± SD. **(G)** Quantification of mCherry-Snf7 punctae of same strain shown in **D** grown to early-log phase in minimal media then shifted to 37°C for six hours. Two trials, n = 40, mean ± SD.

To determine if the Cdc42 spot is a site of Snf7 filament disassembly, we examined the dynamic behavior of the Cdc42 spot and colocalized Snf7 punctae using high time-resolution live-cell imaging. While most of the colocalized Cdc42 and Snf7 spots persisted throughout the duration of imaging sessions, we were able to observe a few instances where a Snf7 spot disassembled while the colocalized Cdc42 signal remained persistent, suggesting that the Cdc42 spot might promote disassembly of Snf7 oligomers *in vivo* (Fig. 4B). To explore the possibility that Vps4 binding toESCRT-III proteins might be facilitated by Cdc42 binding to the ESCRT protein, we performed a GST-pulldown binding assay using purified proteins. We found that Cdc42 interacts with full-length Chm7, the putative “open” form of Chm7 (Chm7^open^), and the “open” form of Snf7 (Snf7^open^), but not to the full-length, “closed” form of Snf7 (Webster et al., 2016; Saksena et al., 2009) (Fig. 4 C). These *in vitro* binding studies suggest that Cdc42 associates with “open,” active Snf7 *in vivo*, and the live-cell imaging suggests that this physical interaction might facilitate disassembly of Snf7 filaments.

To determine whether Cdc42 is required for Snf7 oligomers disassembly *in vivo*, we examined by fluorescence microscopy the number of mCherry-Snf7 punctae in *cdc42-129* mutants. To avoid bias in defining the parameters for punctae selection, we measured the overall frequency distribution of pixel intensities of mCherry-Snf7 signal at the medial focal plane (see methods). Compared to wild-type cells, *cdc42-129* mutants had more high-intensity mCherry-Snf7 pixels than the control. This increase in the amount of large Snf7 assemblies suggests that *cdc42-129* mutants have defects in Snf7 disassembly and turnover (Figs. 4 D, F, G).

Our initial characterization of the Cdc42 spot revealed a curious localization to the site of fission of an ER tubule that bridges the nuclear and cortical ER (Fig. 1 C). The yeast ER network is composed primarily of cortical ER and nuclear ER, with the cytoplasmic space generally devoid of ER tubules (Preuss et al., 1991). Therefore, our observation of the Cdc42 spot at sites of ER tubules undergoing fission prompted us to examine the ER morphology of the *cdc42-129* mutant. We observed that *cdc42-129* mutants exhibit an ER phenotype in which the cytoplasmic space between the nuclear ER and cortical ER is filled with a network of ER tubules emanating from the nuclear ER (Fig. 4 D, E). This phenotype suggests that, in addition to sealing holes in the nuclear envelope, Cdc42 is also involved in maintaining the architecture of the endoplasmic reticulum. Together with the previous section’s findings, these fluorescent microscopy findings support the idea that Cdc42 functions as a regulator of Snf7 disassembly, and that the loss of this regulation results in an increase in Snf7 filaments that cause disorganization of the nuclear envelope and ER.

We have uncovered novel functions of Cdc42 as a regulator of ESCRT-III disassembly during nuclear envelope sealing and in the maintenance of ER morphology. In budding yeast, fission yeast, and mammalian cells, loss of ESCRT-III and LEM function causes persistent holes in the nuclear envelope, ultimately leading to defective nucleo-cytoplasmic partitioning (Vietri et al., 2015; Olmos et al., 2015; Webster et al., 2014; Gu et al., 2017) (Fig. 5 B). The same functional consequence occurs when the ESCRT-LEM complex fails to disassemble. Fission yeast lacking Vps4, and therefore exhibit unrestricted ESCRT function, have persistent fenestrations, peri-nuclear ER stracks (karmellae), and disorganized tubular extensions of the nuclear envelope, culminating in the loss of nucleo-cytoplasmic compartmentalization (Gu et al., 2015) (Fig. 5 C). While the *cdc42-129* mutant exhibits a leaky nucleus phenotype like *snf7*Δ, *chm7*Δ, and *heh1Δheh2*Δ mutants, it also displays increased Snf7 aggregates and sinuous tubular extensions from the nuclear envelope, reminiscent of the disorganized tubular extensions of the nuclear membrane observed in *vps4*Δ fission yeast (Gu et al., 2015). These data suggest that Cdc42 regulates ESCRT disassembly, which is required for both nuclear envelope sealing and maintenance of proper ER architecture in budding yeast. We therefore propose that Cdc42 functions in ESCRT-mediated nuclear envelope sealing as a regulator of ESCRT disassembly, where it appears to function with Vps4 to dismantle the ESCRT-LEM machinery at the nuclear envelope to seal holes in the nuclear envelope as well as maintain ER morphology (Fig. 5 A). Because Cdc42 preferentially binds to the “open” form of Snf7, we speculate that it may act as a cofactor to promote the association of Vps4 and its weakly-binding ESCRT-III partner CHMP4B/Snf7. Taken together, our live-cell imaging, biochemical, and genetic analyses provide evidence for a novel Cdc42 function in nuclear envelope sealing as the first-identified regulator of ESCRT function in this process.

**Figure 5.**
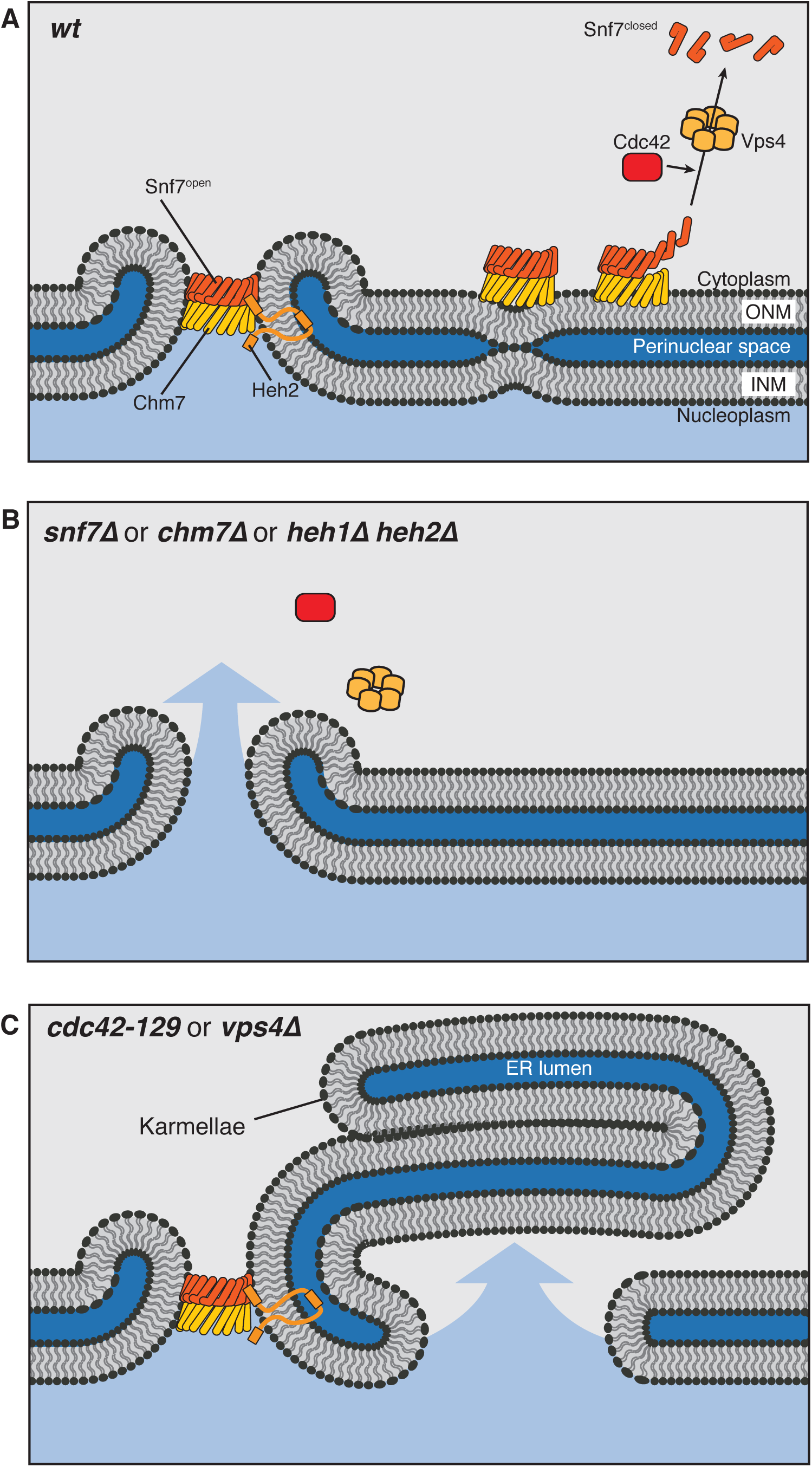
Cdc42, ESCRT-III, and Heh2 mutants share a leaky nucleus phenotype. **(A)** In normal cells, Heh2, Chm7, and Snf7 function at holes in the nuclear envelope to carry out annular fusion. ESCRT-III polymers are disassembled by Vps4 and we propose that Cdc42 is involved in the disassembly step by either directly contributing to Snf7 disassembly, activating Vps4 function, or cooperating with Vps4. **(B)** Mutants lacking components directly involved in annular fusion have holes in their nuclear envelopes left by nuclear fusion and ER fission events. **(C)** Cells lacking Vps4 and normal Cdc42 have unregulated ESCRT activity at the nuclear envelope. This causes the formation of nuclear karmallae and large holes in the nuclear envelope, leading to a defect in proper nucleo-cytoplasmic partitioning.

## Materials and Methods

### Yeast Strain Generation and Growth

All strains and the details of their construction are described in Table 1. Gene knockouts and C-terminal fluorescent protein fusions of endogenous genes were constructed using PCR-based integration of DNA cassettes derived from template plasmids as previously described (Longtine). N-terminal fusions were constructed using PCR-based integration of DNA cassettes generated using overlap extension PCR. The NLS-GFP strain was generated by integrating at the *LEU2* locus an NLS-3xGFP-containing plasmid, which was generously provided by Patrick Lusk (Webster Cell) which is further described in plasmids. The HDEL strain was generated by integrating at the *TPI1* locus a GFP-HDEL containing plasmid, which was generously provided by Laura Lackner/Jodi Nunnari/Randy Schekman (unpublished) and further described in plasmids.

**Table 1.**
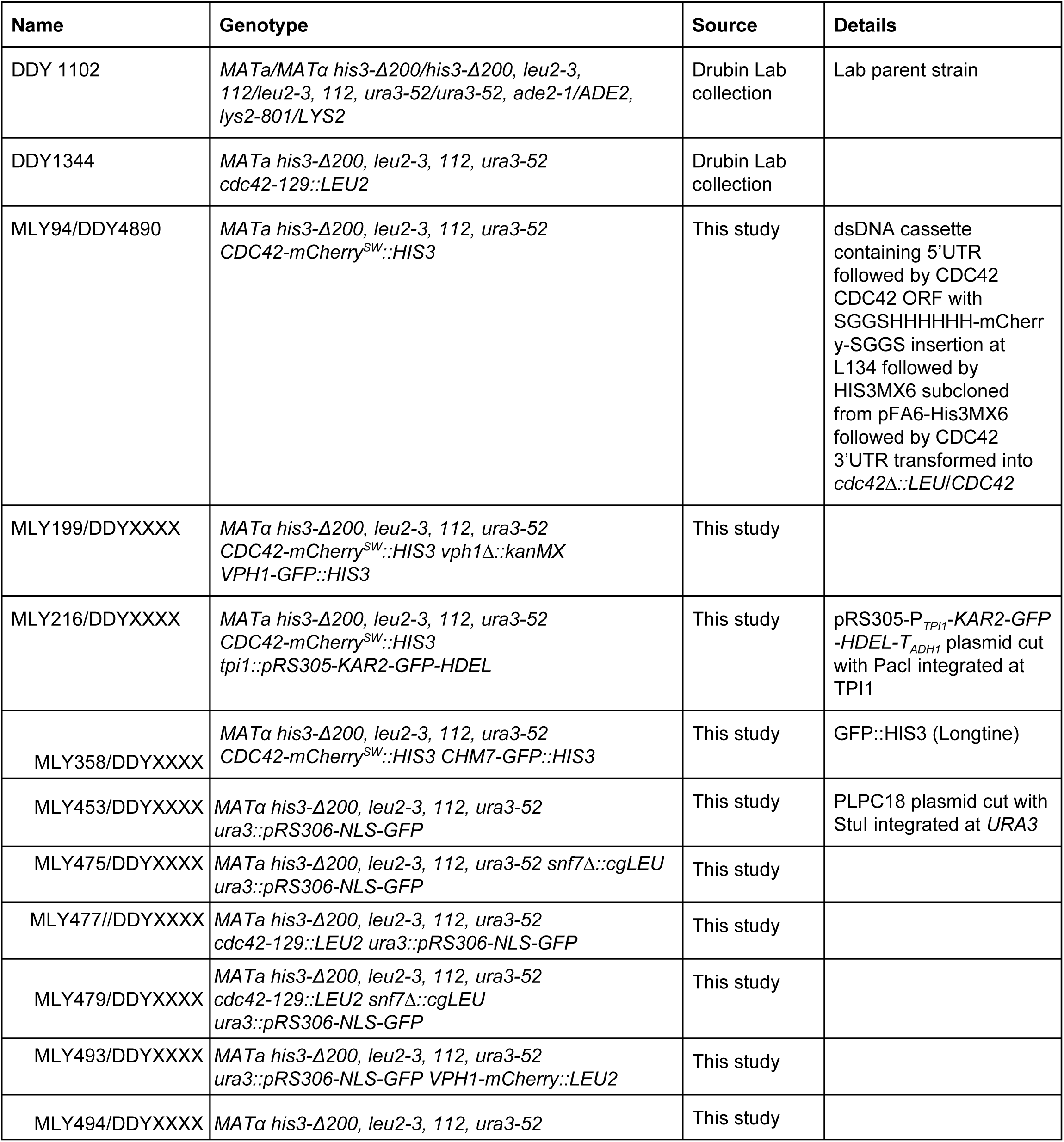
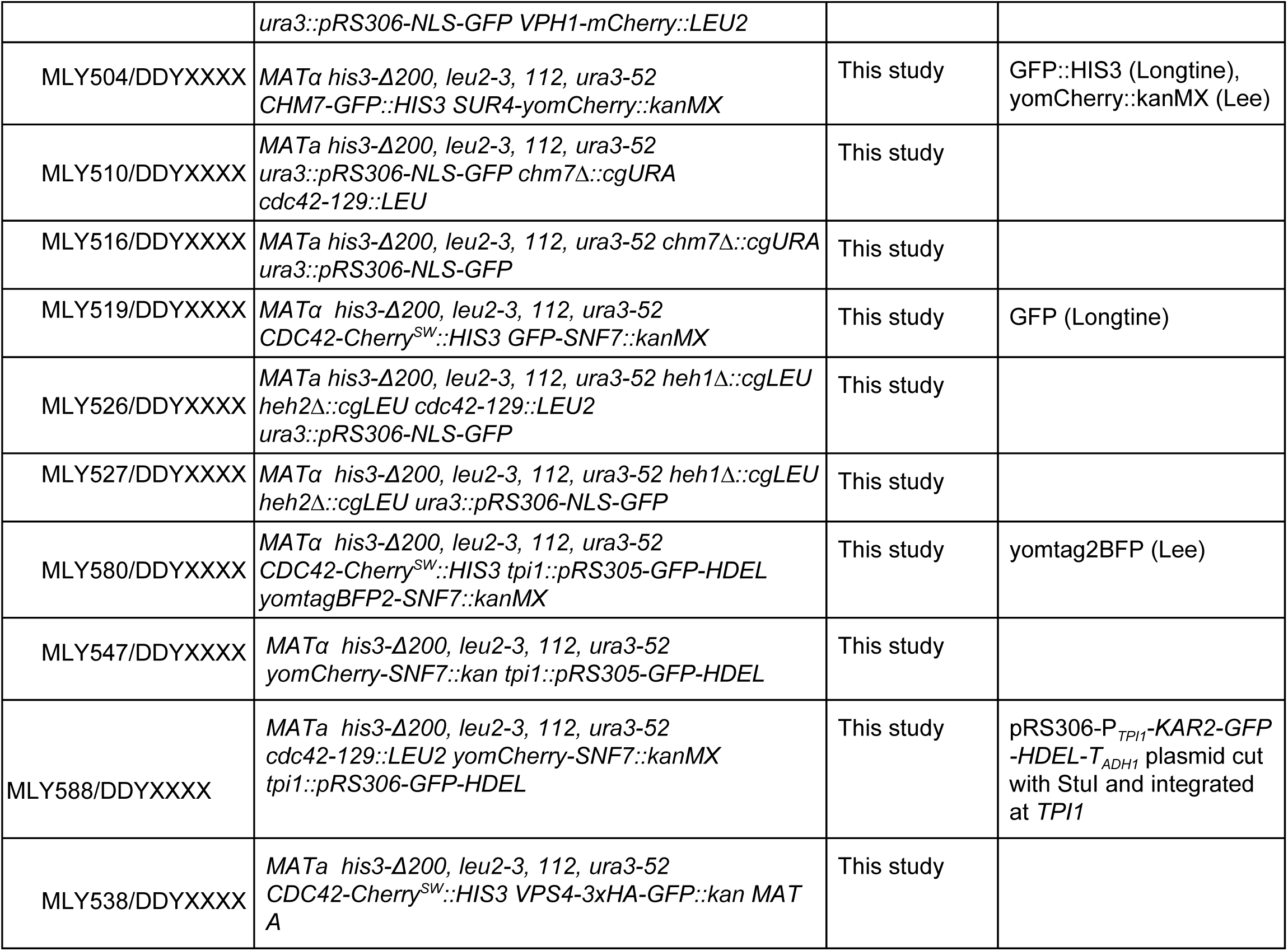
Strains used in this study.

The *CDC42-mCherry^SW^::HIS3* strain was constructed by transforming a *cdc42Δ::LEU/CDC42* diploid strain with a dsDNA cassette (generated using overlap extension PCR) containing the 5’UTR (500nt) of *CDC42* followed by the open reading frame with an SGGSHHHHHH-mCherry-SGGS insertion at Leucine 134, followed by the *HIS3* auxotrophic marker amplified from pFA6a-GFP-HISMX6, followed by 3’UTR (500nt) of *CDC42*. His^+^Leu^-^ diploid transformants were then sporulated and dissected to obtain the *CDC42-mCherry^SW^::HIS* haploid. Strains were propagated using standard techniques (Amberg).

### Plasmids

All plasmids used are listed in Table 2. pFA6a-GFP-HIS3MX6 (Longtine) was used for the PCR integration of C-terminal GFP fusions of endogenous genes. pFA6a-link-yomCherry-KanR (Lee PLoS) was used for the PCR integration of C-terminal mCherry fusions of endogenous genes. Gene deletion cassettes were PCR amplified from *cgLEU* or *cgURA* that were cloned into pBluescript vectors. The NLS-GFP plasmid PLPC18 was provided by Patrick Lusk (Webster Cell) and is a pRS406 plasmid containing the NLS sequence of *HEH2* followed by 3x GFP under the control of the *PHO4* promoter. The *GFP-HDEL::LEU* plasmid used was a gift from Laura Lacker (unpublished), and is a pRS305 plasmid containing the promoter of *TPI1*, followed by the leader sequence of *KAR2* (a.a. 1-52), followed by GFP, followed by HDEL. The URA version of the plasmid was made by subcloning the *TPI1-KAR2-GFP-HDEL* fragment into pRS306. For GST-fusion protein expression, Chm7, Heh2, and Snf7 fragments were cloned into pGEX-4T-1 (GE lifesciences) vector using BamHI and XhoI. The following fragments were used: Heh2^NT^ is amino acids 1-308, Heh2^CT^ is a.a. 566-663, Snf7^open^ is a.a. 1-156, Snf7^FL^ is a.a. 1-240, Chm7^open^ is a.a. 1-369, Chm7^FL^ is a.a. 1-450. For His-Cdc42^Q61L^ protein purification, Full-length Cdc42 with a Q61L point mutation was cloned into a pBH-based vector (Smith) containing a TEV-cleavable N-terminal 6x-His epitope using BamHI and XhoI.

**Table 2.** Plasmids used in this study.

| **Plasmid** | **Use** | **Source** |
| --- | --- | --- |
| pFA6a-GFP(S65T)-HIS3MX6 | C-terminal GFP::HIS integration at endogenous locus | Longtine |
| pFA6a-yomCherry-kanMX | C-terminal mCherry::kan integration at endogenous locus | Lee |
| pFA6a-yomtagBFP2-kanMX | Used to make *BFP-SNF7::kanMX* | Lee |
| pBS-cgLEU | Used to generate deletions | Lab collection |
| pBS-cgURA | Used to generate deletions | Lab collection |
| pRS406-P<sub>PHO8</sub>-NLS(HEH2)-3xGFP (PLPC18) | NLS-GFP integration at *URA3* locus | Webster Cell |
| pRS305-P<sub>TPI1</sub>-KAR2 (a.a. 1-52)-GFP-HDEL-T<sub>ADH1</sub> | Used to make GFP-HDEL integrated at *LEU2* | Laura Lackner (unpublished) |
| pRS306-P<sub>TPI1</sub>-KAR2 (a.a. 1-52)-GFP-HDEL-T<sub>ADH1</sub> | Used to make GFP-HDEL integrated at *URA3* | Laura Lackner (unpublished) |
| pGEX-4T-1-Heh2<sup>1-308</sup> | Protein purification | This study |
| pGEX-4T-1-Heh2<sup>566-663</sup> | Protein purification | This study |
| pGEX-4T-1-Snf7<sup>1-156</sup> | Protein purification | This study |
| pGEX-4T-1-Snf7<sup>1-240</sup> | Protein purification | This study |
| pGEX-4T-1-Chm7<sup>1-369</sup> | Protein purification | This study |
| pGEX-4T-1-Chm7<sup>1-450</sup> | Protein purification | This study |
| pBH-Cdc42<sup>Q61L</sup> | Protein purification | This study (pBH vector from Smith and Prehoda) |

### Fluorescence microscopy

Cells were grown to early-mid log-phase (OD A^600^ 0.1-0.7 reading on Ultrospec 10 Cell Density Meter [Amersham]) from an overnight 1:50 - 1:100 dilution of a log phase culture in imaging media (synthetic minimal media supplemented with adenine, l-histidine, l-leucine, l-lysine, l-methionine, uracil, and 2% glucose). Cells were adhered to 25 mm, 1.5 thickness, circular coverslips (Deckgläser) coated with 0.2 mg/ml concanavalin A.

Epifluorescence imaging was performed on a Nikon Eclipse Ti inverted microscope with a Nikon 100× 1.4-NA Plan Apo VC oil-immersion objective and an Andor Neo 5.5 sCMOS camera. GFP and mCherry fluorescence were excited using a Lumencor Spectra X LED light source with 470/22-nm and 575/25-nm excitation filters, respectively. For GFP-mCherry two-color imaging, channels were acquired sequentially using an FF-596-Di01-25×36 dual-pass dichroic mirror and FF01-641/75-25 dual-pass emission filters (Semrock). The system was controlled with Nikon Elements software and maintained in a room maintained at 23-25**°**C.

Spinning disk confocal imaging was performed on a Nikon Eclipse Ti inverted microscope outfitted with a Nikon 100x 1.45-NA Plan Apo VC or Nikon 60x 1.4-NA Plan Apo VC oil-immersion objective lens, a Yokogawa CSU-X1 spinning disk, and Andor iXon^EM^ DU-897 EMCCD camera. GFP, mCherry, and mtagBFP2 fluorescence were excited using 488-nm, 561-nm, and 405-nm lasers, respectively, and detected with Chroma 535/20-nm, Chroma 605/52-nm, and 470/40-nm emission filters, respectively. The system was controlled with Nikon Elements software and kept in a room maintained at 23-25**°**C.

### GST pulldown assays

Purified GST-Heh2^NT^ (860 nM), purified GST-Heh2^CT^ (900 nM), purified GST-Snf7^FL^ (200 nM), purified GST-Snf7^open^ (280 nM), 1000 µl GST-Chm7^FL^ bacterial lysate, and 1000 µl GST-Chm7^open^ were brought up to 1000 µl in transport buffer (20mM HEPES pH 7.5, 110 mM KOAc, 2mM MgCl2, 1% Triton X-100, 1mM DTT) and incubated with 25 µl glutathione agarose (50:50 v/v slurry in deionized H_2_O) at 4°C for 1 hour. Beads were then washed in 500 µl cold transport buffer three times. Purified His_6_-Cdc42 was then added to the beads at a concentration of 2.1 µM to a final reaction volume of 100 µl, then incubated at 4°C for 1 hr with slight agitation. The beads were then washed in 500 µl transport buffer three times. Proteins were eluted with 13 µl 6x SDS sample buffer (125 mM Tris pH 6.8, 30% glycerol (v/v), 4.1% SDS (w/v), 20 mM DTT)

### SDS-PAGE and western blotting

13 µl of the GST pulldown assay eluates were resolved by SDS-PAGE and either stained with GelCode Blue Stain reagent (Thermo) or transferred onto 0.2 µm nitrocellulose (Amersham Protran) in 25 mM Tris, 120 mM glycine, 20% methanol. Nitrocellulose membranes were then dried for 30 min and blocked in 5% milk in TBS for 30 minutes. Mouse anti-His primary antibody (R&D systems) was used at 1:1000 dilution in TBS, 1% Tween-20, 1% block and applied for 1 hour at room temperature. Blot was washed for 15 minutes in TBS, 1% Tween-20, then IRDye 800CW Goat anti-Mouse secondary antibody (Li-Cor) was used at 1:10,000 dilution in TBS + 1% Tween-20, 0.1% block, and incubated at room temperature for one hour. Blot was then washed for 30 minutes in TBS, 1% Tween-20 and imaged using an Odyssey CLx (Li-Cor).

### Production of recombinant proteins

All proteins were expressed and purified from *E.coli* Rosetta (DE3) Competent Cells (Novagen). Details of individual constructs and their expression plasmids are detailed in the plasmids section. For GST-fusion proteins, cells were grown in LB + ampicillin (100 µg/ml) + chloramphenicol (25 µg/ml) to mid log phase, and expression was induced with 250 µM IPTG for 16 hours at 18°C. Cells were pelleted and resuspended in PBS + Complete Mini EDTA-free protease inhibitor (Roche). Cells were lysed by sonication using a W-385 Sonicator Ultrasonic Processor (Heat Systems - Ultrasonics, Inc), using one minute of 0.5 sec on/0.5 sec off cycling for two cycles. Crude lysates were then adjusted to contain 1% Triton X-100, and clarified by spinning at 25k RPM in a Ti-70 rotor. Clarified lysates were applied to 1mL glutathione agarose (Sigma), incubated at 4°C for an hour, washed with 150 mL PBS + 1% Triton X-100, then eluted with 30 mM reduced glutathione in 50 mM Tris pH 9.5. GST-fusion protein eluates were then dialyzed overnight at 4°C in 25 mM Tris pH 7.5, 150 mM NaCl, 1 mM DTT using a Slide-A-Lyzer 10 kDa MWCO cutoff dialysis cassette (Thermo). The following day, dialysates were concentrated over a 10k, 30k, or 50k Amicon Ultra-4 Centrifugal filter (Millipore). Concentrated, purified protein was aliquoted, snap frozen in liquid nitrogen, and stored at −80°C.

For His_6_-Cdc42 purification, protein expression was performed as described for GST-fusion proteins. Harvested cells were pelleted and resuspended in PBS + 20 mM Imidazole. Cell lysis and lysate clarification were performed as described above. Clarified lysate was then incubated with 1 mL Ni-NTA agarose (Invitrogen) for one hour at 4°C. Beads were then washed in 150 mL PBS + 20 mM imidazole and protein was eluted with 250 mM imidazole. Eluate was then desalted over a PD-10 column (GE lifesciences) into 50 mM Tris pH 8.0, 150 mM NaCl and polished over a mono Q 5/50 GL anion exchange column (GE Lifesciences). Bound proteins were eluted off over 20 column volumes to 35% elution buffer (50 mM Tris pH 8.0, 1 M NaCl). Eulates were dialyzed overnight against 25 mM Tris pH 7.5, 150 mM NaCl, 1 mM DTT using a Slide-A-Lyzer 10 kDa MWCO dialysis cassette (Thermo). The following morning, the purified protein was concentrated over a 10k Amicon Ultra-4 Centrifugal filter (Millipore) then immediately aliquoted, snap frozen in liquid nitrogen, and stored at −80 °C until further use.

The concentrations of purified proteins were measured by SDS-PAGE using GelCode Blue Stain Reagent (Thermo) to visualize protein bands. The quantification of coomassie-stained bands was determined using a Li-Cor Odyssey CLx and its accompanying software. Ultra pure Bovine Serum Albumin supplied in the BCA Protein Assay kit (Thermo) was used to generate a standard curve.

### Imaging processing and analysis

All micrographs were acquired using Nikon Elements imaging software. All images were processed using FIJI/ImageJ. Kymographs in Figures 1c, 1d, 2a-c were generated from maximum intensity projections of 2µm at 0.5µm steps using a one-pixel line drawn across the Cdc42 spot. The “Plot profile” tool was used to obtain intensity values as a function of distance, which was then exported and re-plotted in PRISM (Graphpad) as values normalized to the maximum value.

To measure the nuclear accumulation of NLS-GFP, we measured the GFP fluorescence intensity of a 0.5 µm^2^ area in the nucleus and a 0.5 µm^2^ area in the cytoplasm from maximum intensity projection images of 11 µm at 0.5 µm steps. The background was averaged and subtracted from NLS-GFP values, and the nuclear accumulation was plotted as a mean ratio of nuclear and cytoplasmic measurements. Individual cell measurements are shown in EXT DATA FIGURE 4, where the three individual trials for each mutant is represented by a circle, triangle, or square, and open shapes represent the wildtype control in the same field of view. The averages of each trial of each mutant was then normalized to the average of the wildtype control to generate normalized average NLS-GFP ratios shown in FIGURE 4e.

Frequency distributions shown in Figure 5c, d were generated from measurements taken from 20 cells per trial for two trials using 16-bit grayscale images. A single focal plane at the thickest part of the cell was analyzed instead of intensity projections because of the large variation in cell size of the mutant strain. A boundary was drawn around the edge of the cell’s medial focal plane (cell cortex boundary determined by cortical GFP-HDEL in the green channel) using the “freehand selections” tool in ImageJ, then a histogram of pixel intensities was generated using the “histogram” function where the bin-width was set to one. The area in µm^2^ of the cell was then measured using the “Measure” function, and this value was used to normalize pixel counts to area. The frequency distribution values were then re-plotted using PRISM (Graphpad) software. The mean is represented by the thick line, and the standard deviation is represented as 1pt vertical lines.

To determine the ER area/cell area ratio for each cell, ImageJ was used to analyze 16-bit confocal spinning disk micrographs of a single focal plane at each cell’s thickest portion. The “freehand selection” tool was used to outline the boundary of each cell. Then, the boundaried pixels were thresholded using the “moment” algorithm (Tsai) and converted to binary. ER area was calculated as percentage of signal pixels (255 value pixels) to total pixels (sum of 0 value pixels and 255 value pixels).

## Supplemental Materials

Fig. S1 shows growth assay of the Cdc42-mCherry^SW^ strain, as well as additional examples of the Cdc42 focus in the mother, bud, and mother and bud. It also shows a montage from video 1. It also contains a cartoon illustration of the endoplasmic reticulum and nuclear membrane architecture. Fig. S2 shows additional localization fluorescence microscopy micrographs of the Cdc42 spot at the vacuole and ER membranes. It also shows a montage taken from video 4of the Cdc42 spot at the vertices of fusing vacuoles. Fig. S3 shows the cytoplasmic:nucleus NLS-GFP measurements obtained for each cell of evert trials of the reported averages shown in Fig. 3 E.

## Acknowledgements

We thank Patrick Lusk and Laura Lackner for generously providing plasmids. We thank Ross Pedersen and James Hurley for critically evaluating the manuscript, and members of the Drubin/Barnes laboratory for constant informal input. We thank H. Aaron for her microscopy training and assistance. Spinning disc confocal microscopy was conducted at the University of California, Berkeley, Cancer Research Laboratory Molecular Imaging center, supported by the Gordon and Betty Moore Foundation. This work was supported by the National Institutes of Health grant R35GM118149 to D.G. Drubin. The authors declare no competing financial interests.

## Author Contributions

M.S. Lu and D.G. Drubin conceived of the experiments. M.S. Lu generated the reagents, performed the experiments, and analyzed the data. M.S. Lu and D.G. Drubin wrote the manuscript. D.G. Drubin secured funding.

## Supplemental Information

### Supplemental Figures

**Figure S1.**
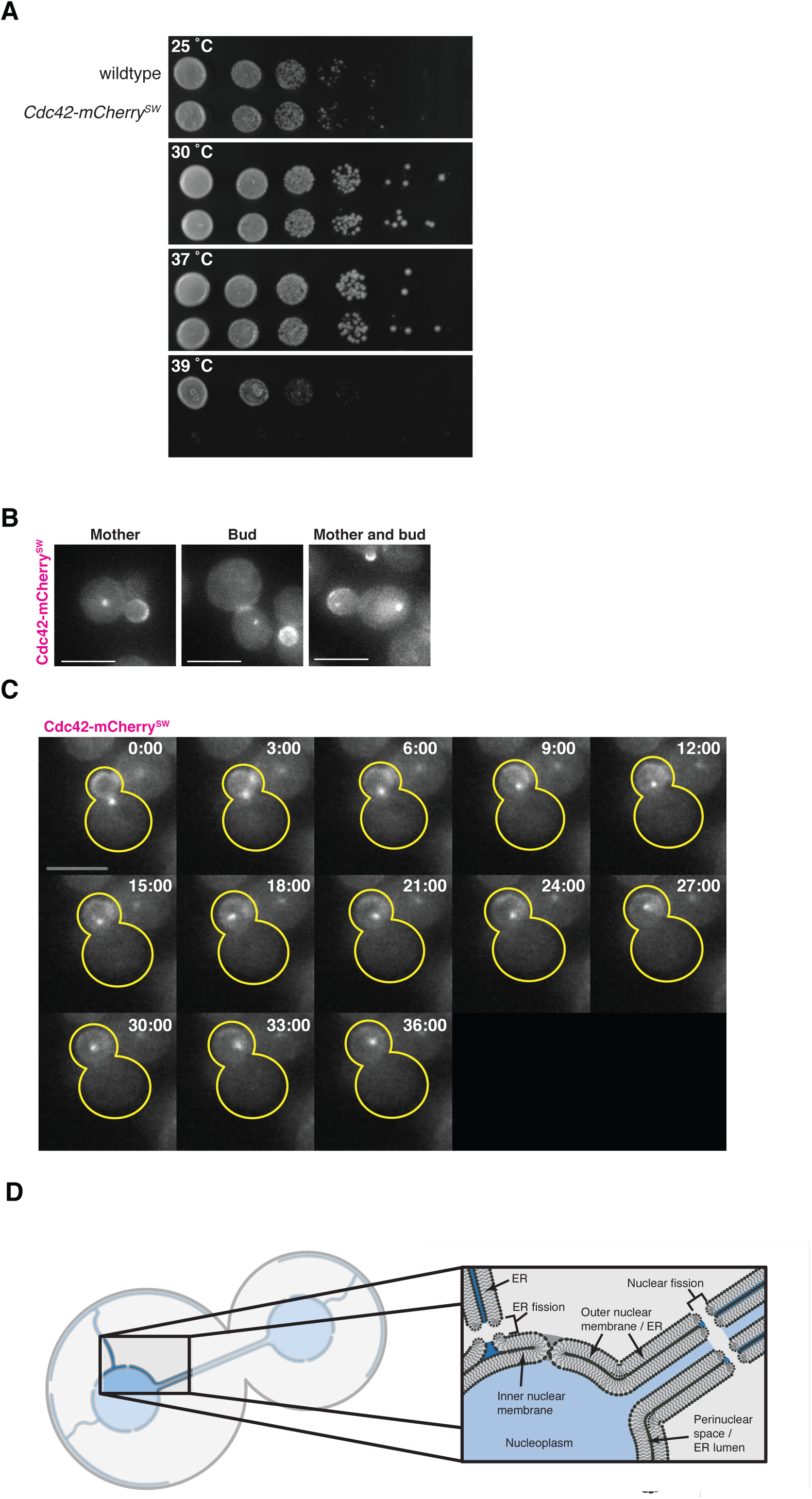
A functional internal fluorescent protein fusion of Cdc42 reveals a novel subcellular localization. **(A)** Cell growth of indicated yeast strains was compared by spotting serial dilutions of liquid cultures on plates at 25°C, 30°C, 37°C, or 39°C. **(B)** Micrographs of three separate cells expressing endogenous Cdc42-mCherry^SW^ showing Cdc42 spot in the mother, the bud, or mother and bud. **(C)** Montage of confocal spinning disk movie stills of cells endogenously expressing Cdc42-mCherry^SW^ showing Cdc42 spot segregating from mother cell into bud. Time is in minutes. All cells in micrographs of this figure were imaged during log phase in minimal imaging media at 23-25 °C. All scale bars, 5µm. 2µm maximum intensity projections at 0.5µm steps. **(D)** Illustration depicting the organization of membranes and lumen spaces of the nuclear envelope and endoplasmic reticulum. The perinuclear space is continuous with the ER lumen (dark blue) because the outer nuclear membrane is continuous with the ER membrane. Nuclear fission events introduce a break in the neck of the dividing nucleus to expose the nucleoplasm (light blue) to the cytosolic space.

**Figure S2.**
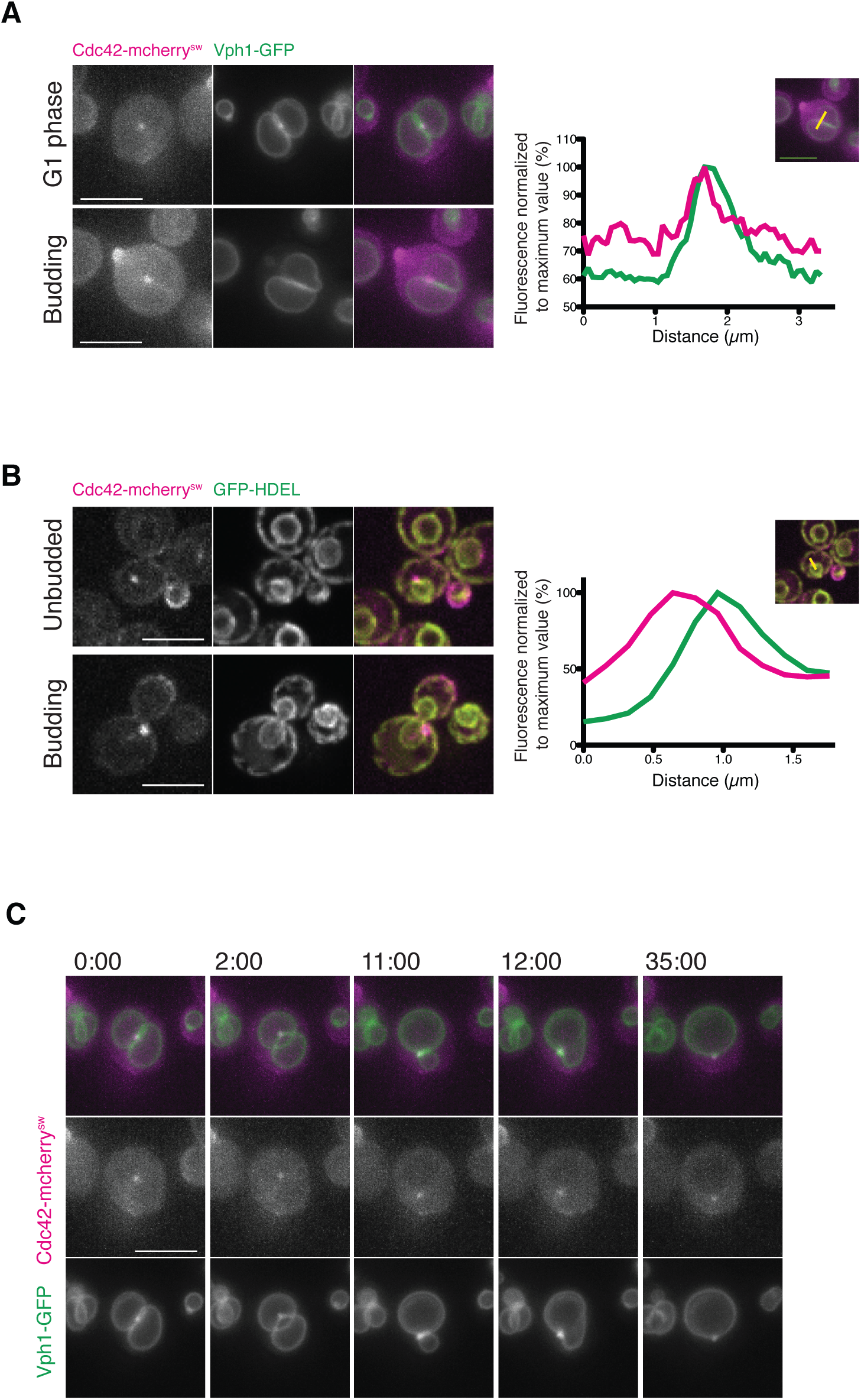
The Cdc42 spot is localized to vacuolar and ER membranes, and vertices of fusing vacuoles. **(A)** Epifluorescence still images of cells endogenously expressing Cdc42-mCherry^SW^ (magenta) and Vph1-GFP (green) in unbudded (top panel) or budded cell (lower panel). Plot of normalized fluorescence intensity profiles of Vph1-GFP (green) and Cdc42-mCherry^SW^ (magenta) along yellow line shown in inset image. **(B)** Epifluorescence still images of cells endogenously expressing Cdc42-mCherry^SW^ (magenta) and GFP-HDEL (green) in unbudded (top panel) or budded cell (lower panel). Plot of normalized fluorescence intensity profiles of GFP-HDEL (green) and Cdc42-mCherry^SW^ (magenta) along yellow line shown in inset image. All images are 2µm maximum intensity projections at 0.5µm steps. All scale bars 5µm. **(C)** Montage of stills from epifluorescence movies of cells expressing endogenous Cdc42-mCherry^SW^ (magenta) and Vph1-GFP (green). Cells were imaged were in log phase growth in minimal imaging media at 23-25 °C. Scale bar, 5µm. 2µm maximum intensity projections at 0.5µm steps.

**Figure S3.**
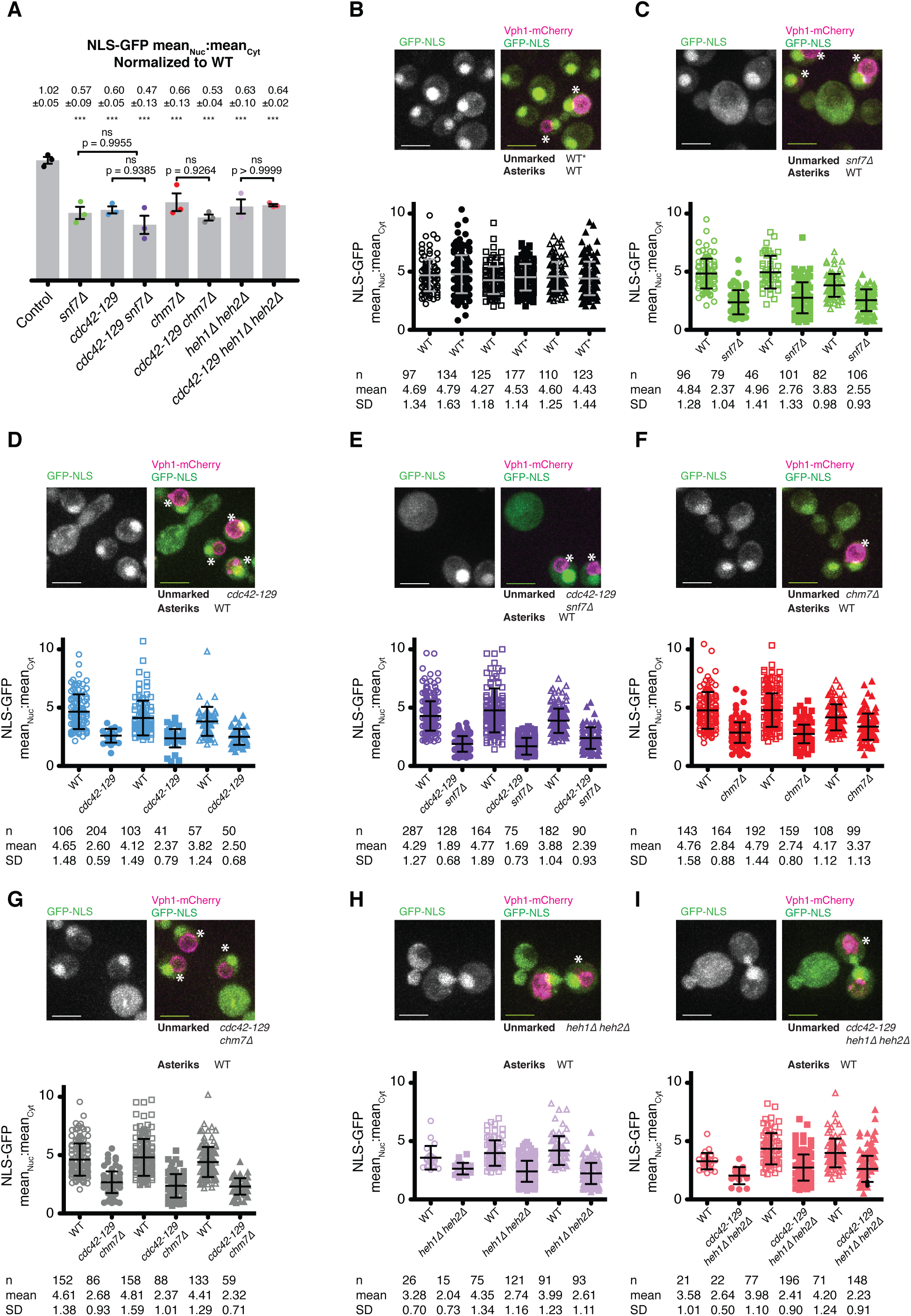
Cdc42 functions in same NE sealing pathway as ESCRT and LEM proteins. **(A)** same quantification shown in Figure 3 E, but with Mean ± SD figures reported on top of bar graph. **(B)** WT control. Confocal spinning disk micrograph of control strain and experimental strain in the same field of view, where cells denoted with asterisks are the control WT strain. Left micrograph shows NLS-GFP signal only, whereas right micrograph also shows Vph1-mCherry signal, which was used to identify WT control cells. Scatter plot shows individual measurements of mean nucleus:cytosol GFP measurement from all trials, where circles, triangles, and squares represent different trials. Open shapes indicate measurements from control strains and closed shapes from mutant strain. n, mean, and standard deviation reported beneath plot for each trial for control and mutant. **(C)** *snf7*Δ. **(D)** *cdc42-129*. **(E)** *cdc42-129 snf7*Δ. **(F)** *chm7*Δ. **(G)** *cdc42-129 chm7*Δ. **(H)** *heh1 heh2*. **(I)** *cdc42-129 heh1 heh2*.

### Supplemental Videos

**Video 1.** Cdc42 (magenta) spot segregates to bud during division. ER (green) shown to clarify cell boundary. Scale bar 5µm. Time in minutes.

**Video 2.** Cdc42 (magenta) spot localizes to the base of an ER (green) tubule undergoing fission. Scale bar 5µm. Time in minutes.

**Video 3.** Cdc42 (magenta) spot localizes to the base of the neck of the nuclear envelope undergoing fission. Scale bar 5µm. Time in minutes.

**Video 4.** Cdc42 (magenta) spot localizes to vertices of fusing vacuole (green) fragments. Scale bar 5µm. Time in minutes.

**Video 5.** Cdc42 (red) and Snf7 (blue) colocalized spot migrate to the base of the neck of the nuclear envelope (green) undergoing fission. Scale bar 5µm. Time in minutes.

